# Heterogeneities in intrinsic excitability and frequency-dependent response properties of granule cells across the blades of the rat dentate gyrus

**DOI:** 10.1101/701342

**Authors:** Poonam Mishra, Rishikesh Narayanan

## Abstract

The dentate gyrus (DG), the input gate to the hippocampus proper, is anatomically segregated into three different sectors, namely the suprapyramidal blade, the crest region and the infrapyramidal blade. Although there are well-established differences between these sectors in terms of neuronal morphology, connectivity patterns and activity levels, differences in electrophysiological properties of granule cells within these sectors have remained unexplored. Here, employing somatic whole-cell patch-clamp recordings from the rat DG, we demonstrate that granule cells in these sectors manifest considerable heterogeneities in their intrinsic excitability, temporal summation, action potential characteristics and frequency-dependent response properties. Across sectors, these neurons showed positive temporal summation of their responses to inputs mimicking excitatory postsynaptic currents, and showed little to no sag in their voltage responses to pulse currents. Consistently, the impedance amplitude profile manifested low-pass characteristics and the impedance phase profile lacked positive phase values at all measured frequencies, voltages and for all sectors. Granule cells in all sectors exhibited class I excitability, with broadly linear firing rate profiles, and granule cells in the crest region fired significantly less action potentials compared to those in the infrapyramidal blade. Finally, we found weak pairwise correlations across the 18 different measurements obtained individually from each of the three sectors, providing evidence that these measurements are indeed reporting distinct aspects of neuronal physiology. Together, our analyses show that granule cells act as integrators of afferent information, and emphasize the need to account for the considerable physiological heterogeneities in assessing their roles in information encoding and processing.

## INTRODUCTION

The dentate gyrus (DG), the input gate to the mammalian hippocampus proper (Amaral et al. 2007; Andersen et al. 2006), has been implicated in spatial navigation, response decorrelation, pattern separation and engram formation. Granule cells are the prominent neuronal subtype within the DG, and have been studied extensively from the perspective of their intrinsic response properties, plasticity profiles, *in vivo* response properties, their role as engram cells, their sparse connectivity and sparse firing characteristics, and neurogenesis (Aimone et al. 2014; Amaral et al. 2007; Bakker et al. 2008; Bliss and Lomo 1973; Danielson et al. 2017; Diamantaki et al. 2016; GoodSmith et al. 2017; Heigele et al. 2016; Kropff et al. 2015; Leutgeb et al. 2007; Li et al. 2017; McHugh et al. 2007; Mishra and Narayanan 2019; Neunuebel and Knierim 2014; Sahay et al. 2011; Senzai and Buzsaki 2017; Tonegawa et al. 2018). Electrophysiological recordings from granule cells have characterized their response characteristics, including important differences between mature and immature cell excitability (Fricke and Prince 1984; Krueppel et al. 2011; Liu et al. 1996; Mody et al. 1992; Pedroni et al. 2014; Schmidt-Hieber et al. 2004; 2007; Staley et al. 1992; van Praag et al. 2002). The dentate gyrus, within each location along its dorso-ventral span, is anatomically segregated into three different sectors: the suprapyramidal blade, the crest region and the infrapyramidal blade (Amaral et al. 2007). There are several well-established differences across these three sectors (Amaral et al. 2007), including morphological differences (Claiborne et al. 1990; Desmond and Levy 1985; 1982; Gallitano et al. 2016; Green and Juraska 1985; Schneider et al. 2014), connectivity patterns (Claiborne et al. 1986), the ratio of basket cells to granule cells (Seress and Pokorny 1981) and activity levels (Chawla et al. 2005; Marrone et al. 2012a; Marrone et al. 2012b; Ramirez-Amaya et al. 2013; Ramirez-Amaya et al. 2006; Ramirez-Amaya et al. 2005; Satvat et al. 2012). Despite this, differences in electrophysiological properties of granule cells present within these sectors have surprisingly remained unexplored. In addition, as the DG is present within an oscillatory network (Bland 1986; Buzsaki 2002; Colgin 2013; 2016; Sainsbury and Bland 1981; Winson 1978; 1974), it is important that neuronal response properties are assessed in a frequency-dependent manner, rather than being confined to steady-state measures of subthreshold excitability (Cole 1941; Cole and Baker 1941a; b; Das et al. 2017; Hu et al. 2002; Hutcheon and Yarom 2000; Krueppel et al. 2011; Mauro et al. 1970; Narayanan and Johnston 2008; 2007; Pike et al. 2000; Schmidt-Hieber et al. 2007; Stegen et al. 2012). The frequency-dependent response characteristics of DG granule cells, however, have also not been systematically characterized across these three DG sectors.

To fill these lacunae, in this study, we performed patch-clamp electrophysiological recordings of granule cells from the three sectors of the rat DG, and systematically measured their electrophysiological characteristics. We show that the granule cells in these different DG sectors manifest considerable heterogeneities in their intrinsic excitability, temporal summation, action potential characteristics and frequency-dependent response properties. We found that the subthreshold excitability measures were dependent on membrane voltage, with significant hyperpolarization-induced reduction in the gain of granule cells across all sectors. Across sectors, these neurons showed positive temporal summation of their responses to current injections that mimicked excitatory postsynaptic currents, and showed little to no sag in their voltage responses to hyperpolarizing or depolarizing pulse current injections. Consistently, the impedance amplitude profile manifested low-pass characteristics and the impedance phase profile distinctly lacked positive phase values at all measured frequencies, voltages and for all DG sectors.

Granule cells across the three DG sectors exhibited class I excitability where they were able to fire action potentials at arbitrarily low firing rates, with broadly linear profiles of firing rate *vs*. current injection (*f–I*) curves. Together, the low-pass frequency-response characteristics, the lack of positive impedance phase, and the linear *f–I* curve showing class I excitability point to DG neurons across all these sectors acting as integrators of afferent information. We found no significant differences in subthreshold response properties of these neurons across the three DG sectors. However, we found that granule cells in the crest region fired less action potentials, in response to suprathreshold current injections, when compared with their counterparts in the infrapyramidal blade. Finally, we assessed correlations across the 18 different sub- and supra-threshold measurements for each of the three DG sectors, and found a large number of measurement pairs showing weak pairwise correlations. This large subset of uncorrelated measurements suggested that the set of measurements employed here in characterizing DG granule cells are assessing distinct aspects of their physiology. Together, our analyses show that dentate gyrus neurons act as integrators of afferent information, and emphasize the need to account for the considerable heterogeneities inherent to this population of neurons in assessing their physiology, including engram formation and their ability to perform channel and pattern decorrelation.

## MATERIALS AND METHODS

### Ethical approval

All experiments reported in this study were performed in strict adherence to the protocols cleared by the Institute Animal Ethics Committee (IAEC) of the Indian Institute of Science, Bangalore. Experimental procedures were similar to previously established protocols (Ashhad et al. 2015; Ashhad and Narayanan 2016; Das and Narayanan 2017; Narayanan et al. 2010; Narayanan and Johnston 2008; 2007; Rathour et al. 2016) and are detailed below.

### Slice preparation for *in-vitro* patch clamp recording

Male Sprague-Dawley rats of 5- to 8-week age were anesthetized by intraperitoneal injection of a ketamine-xylazine mixture. After onset of deep anesthesia, assessed by cessation of toe-pinch reflex, transcardial perfusion of ice-cold cutting solution was performed. The cutting solution contained 2.5 mM KCl, 1.25 mM NaH_2_PO_4_, 25 mM NaHCO_3_, 0.5 mM CaCl_2_, 7 mM MgCl_2_, 7 mM dextrose, 3 mM sodium pyruvate, and 200 mM sucrose (pH 7.3, ∼300 mOsm) saturated with 95% O_2_ and 5% CO_2_. Thereafter, the brain was removed quickly and 350-µm thick near-horizontal slices were prepared from middle hippocampi (Bregma, –6.5 mm to –5.1 mm), using a vibrating blade microtome (Leica Vibratome), while submerged in ice-cold cutting solution saturated with 95% O_2_ and 5% CO_2_. The slices were then incubated for 10-15 mins at 34° C in a chamber containing the holding solution (pH 7.3, ∼300 mOsm) with the composition of: 125 mM NaCl, 2.5 mM KCl, 1.25 mM NaH_2_PO_4_, 25 mM NaHCO_3_, 2 mM CaCl_2_, 2 mM MgCl_2_, 10 mM dextrose, 3 mM sodium pyruvate saturated with 95% O_2_ and 5% CO_2_. Thereafter the slices were kept in a holding chamber at room temperature for at least 45 min before the start of recordings.

### Electrophysiology: Whole-cell current-clamp recording

For electrophysiological recordings, slices were transferred to the recording chamber and continuously perfused with carbogenated artificial cerebrospinal fluid (ACSF/extracellular recording solution) at a flow rate of 2–3 mL/min. All neuronal recordings were performed under current-clamp configuration at physiological temperatures (32–35° C), achieved through an inline heater that was part of a closed-loop temperature control system (Harvard Apparatus). The carbogenated ACSF contained 125 mM NaCl, 3 mM KCl, 1.25 mM NaH_2_PO_4_, 25 mM NaHCO_3_, 2 mM CaCl_2_, 1 mM MgCl_2_, 10 mM dextrose (pH 7.3; ∼300 mOsm). Slices were first visualized under a 10× objective lens to locate the granule cell layer of the dentate gyrus and then 63× water immersion objective lens was employed to perform patch-clamp recordings from DG granule cells, through a Dodt contrast microscope (Carl Zeiss Axioexaminer). Whole-cell current-clamp recordings were performed from visually identified dentate gyrus granule cell somata, using Dagan BVC-700A amplifiers. Borosilicate glass electrodes with electrode tip resistance between 2–6 MΩ (more often electrodes with ∼4 MΩ tip resistance were used) were pulled (P-97 Flaming/Brown micropipette puller; Sutter) from thick glass capillaries (1.5 mm outer diameter and 0.86 mm inner diameter; Sutter) and used for patch-clamp recordings. The pipette solution contained 120 mM K-gluconate, 20 mM KCl, 10 mM Hepes, 4 mM NaCl, 4 mM Mg-ATP, 0.3 mM Na-GTP, and 7 mM K2-phosphocreatine (pH 7.3 adjusted with KOH, osmolarity ∼300 mOsm).

Series resistance was monitored and compensated online using the bridge-balance circuit of the amplifier. Experiments were discarded only if the initial resting membrane potential was more depolarized than –60 mV and if series resistance rose above 30 MΩ, or if there were fluctuations in temperature and ACSF flow rate during the course of the experiment. Unless otherwise stated, experiments were performed at the initial resting membrane potential (reported here as *V*_RMP_) of the cell. Voltages have not been corrected for the liquid junction potential, which was experimentally measured to be ∼8 mV.

### Sub-threshold measurements

We characterized DG granule neurons using several electrophysiological measurements obtained through pulse-current and frequency-dependent current injections (Ashhad et al. 2015; Ashhad and Narayanan 2016; Das and Narayanan 2017; Malik et al. 2016; Mishra and Narayanan 2019; Narayanan et al. 2010; Narayanan and Johnston 2008; 2007; Rathour et al. 2016). Input resistance (*R*_in_) was measured as the slope of a linear fit to the steady-state *V-I* plot obtained by injecting subthreshold current pulses of amplitudes spanning –50 to +50 pA, in steps of 10 pA (Fig. 1*A*). Owing to very high input resistance of many cells, and to avoid spike generation for positive current injections, we also performed recordings in response –25 to +25 pA current injection, in steps of 5 pA. To assess temporal summation, five α-excitatory postsynaptic potentials (α-EPSPs) with 50 ms interval were evoked by current injections of the form *I*_α_ =*I*_max_ *t* exp (–α*t*), with α = 0.1 ms^−1^ (Fig. 1*B*). Temporal summation ratio (*S*_α_) in this train of five EPSPs was computed as *E*_last_/*E*_first_, where *E*_last_ and *E*_first_ were the amplitudes of last and first EPSPs in the train, respectively. Percentage sag was measured from the voltage response of the cell to a hyperpolarizing current pulse of 100 pA and was defined as 100 (1–*V*_ss_/*V*_peak_), where *V*_ss_ and *V*_peak_ depicted the steady-state and peak voltage deflection from *V*_RMP_, respectively.

**Figure 1.**
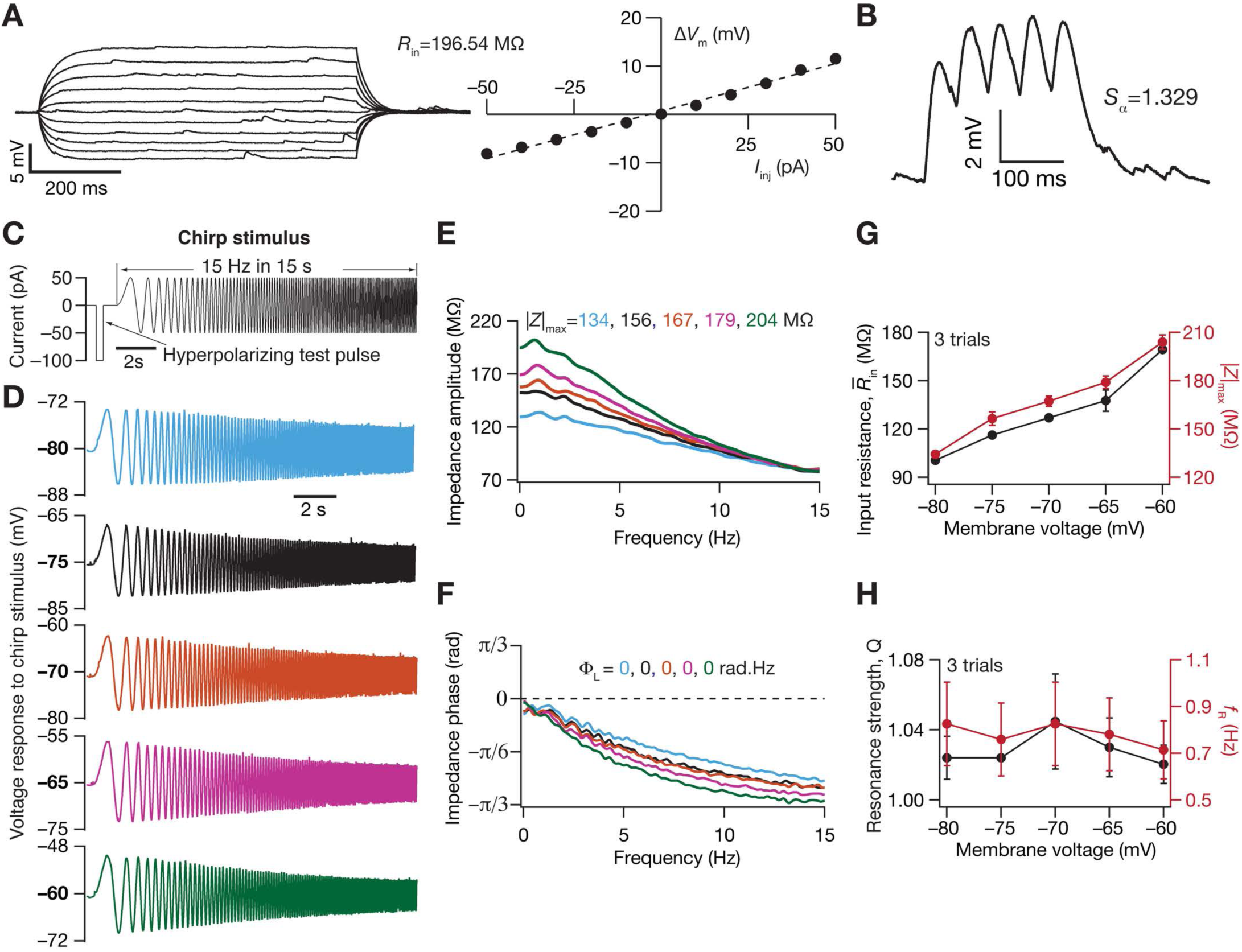
Electrophysiological protocols and measurements employed in the characterization of subthreshold excitability and frequency-dependent response properties of dentate gyrus granule cells. *A*: Voltage responses (*left*) of an example neuron to 700 ms current pulses of amplitude varying from –50 pA to +50 pA (in steps of 10 pA). Input resistance (*R*_in_) was calculated as the slope of the plot (*right*) depicting steady-state voltage response as a function of the injected current amplitude. *B*: Voltage response of the example neuron to 5 alpha-current injections arriving at 20 Hz, depicting temporal summation. Temporal summation ratio (*S*_α_) was computed as the ratio of the amplitude of the fifth response to that of the first. *C*: Chirp stimulus employed for assessing frequency-dependent response properties of the dentate gyrus granule cells. The chirp stimulus employed here was a sinusoidal current of constant amplitude, with frequency varying linearly from 0–15 Hz over a 15 s period. A large 100-pA hyperpolarizing current pulse was provided before the chirp current to estimate input resistance (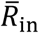) and to observe and correct series resistance changes through the course of the experiment. *D*: Voltage responses of the example neuron to the chirp current at different voltages. The color code for voltages here continue to panels *E* and *F*. *E–F*: Impedance amplitude (*E*) and phase (*F*) profiles computed from the current stimulus shown in panel *C* and the voltage responses shown in panel *D*. |*Z*|_max_ represents the maximum impedance amplitude and Φ_L_ represents the total inductive phase. *G*: Input resistance estimate (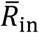) from a single hyperpolarizing pulse (panel *C*) and |*Z*|_max_ (panel *D*) plotted as functions of the membrane voltage at which the chirp stimulus responses were measured. *H*: Resonance strength (*Q*) and resonance frequency (*f*_R_) plotted as functions of the membrane voltage at which the chirp stimulus responses were measured.

The chirp stimulus (Fig. 1*C*) used for characterizing the impedance amplitude (ZAP) and phase (ZPP) profiles was a sinusoidal current of constant amplitude below firing threshold, with its frequency linearly spanning 0–15 Hz in 15 s (*Chirp15*). The voltage response of the neuron was recorded for *Chirp15* current stimulus injection at *V*_RMP_, and for various different voltage values (Fig. 1*D*). The magnitude of the ratio of the Fourier transform of the voltage response to the Fourier transform of the *Chirp15* stimulus formed the impedance amplitude profile (Fig, 1*E*). The frequency at which the impedance amplitude reached its maximum was the resonance frequency (*f*_R_). Resonance strength (*Q*) was measured as the ratio of the maximum impedance amplitude to the impedance amplitude at 0.5 Hz (Hu et al. 2002). Total inductive phase (Φ_L_) was defined as the area under the inductive part of the ZPP (Narayanan and Johnston 2008).

To characterize these subthreshold physiological measurements (see Table 1) of granule cells, recordings were performed at *V*_RMP_ from the three major well-defined sectors of dentate gyrus (Fig. 2): suprapyramidal blade, crest region and infrapyramidal blade (Amaral et al. 2007). The recordings were uniformly distributed within the granule cell layer, across deep, superficial and medial regions (along the hilus-molecular layer axis), of these sectors. Images of cell location were stored for post-facto classification into one of three granule-cell sectors. The characterization protocol to measure sub-threshold measurements was repeated for a range of membrane voltages in a subset of cells, to assess the dependence of these measurements on membrane voltage (Fig. 1*D–H*; Fig. 3).

**Figure 2.**
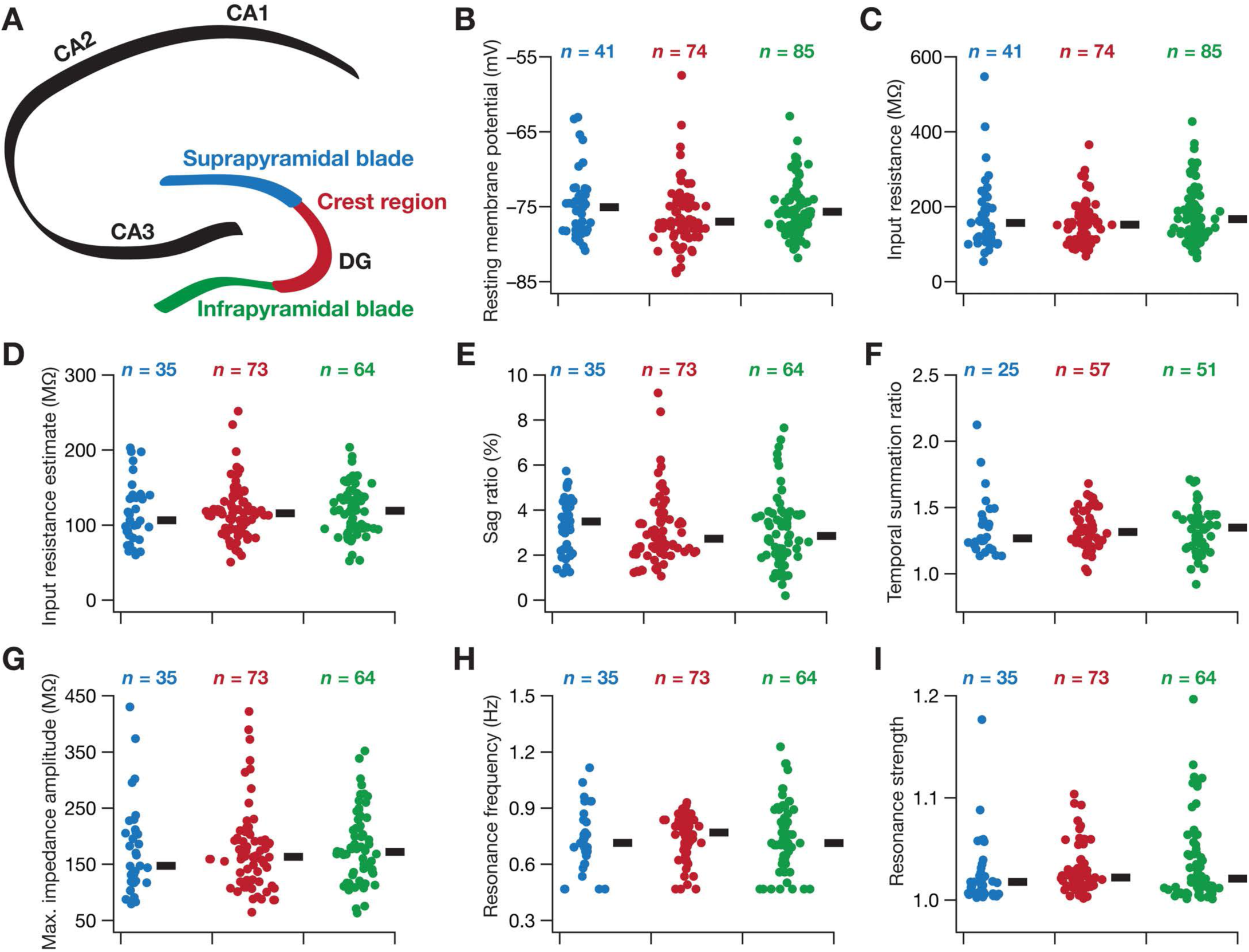
Heterogeneity in subthreshold response properties of granule cells across the blades of the dentate gyrus. *A*: Schematic of the hippocampus proper, showing the different CA (cornu ammonis) subregions (CA1, CA2 and CA3) and the three sectors of the dentate gyrus (infrapyramidal blade, crest region and suprapyramidal blade). The color codes associated with the three DG sectors apply for panels *B*–*I*. *B*–*I*: Beeswarm plots depicting the heterogeneous subthreshold measurements from the three DG sectors. The black rectangles to the right of each beeswarm plot represent the median for the specified population. All measurements depicted in this figure were obtained through current injections into a cell resting at *V*_RMP_. None of the eight subthreshold measurements were significantly different across the three sectors (Kruskal Wallis test).

**Figure 3.**
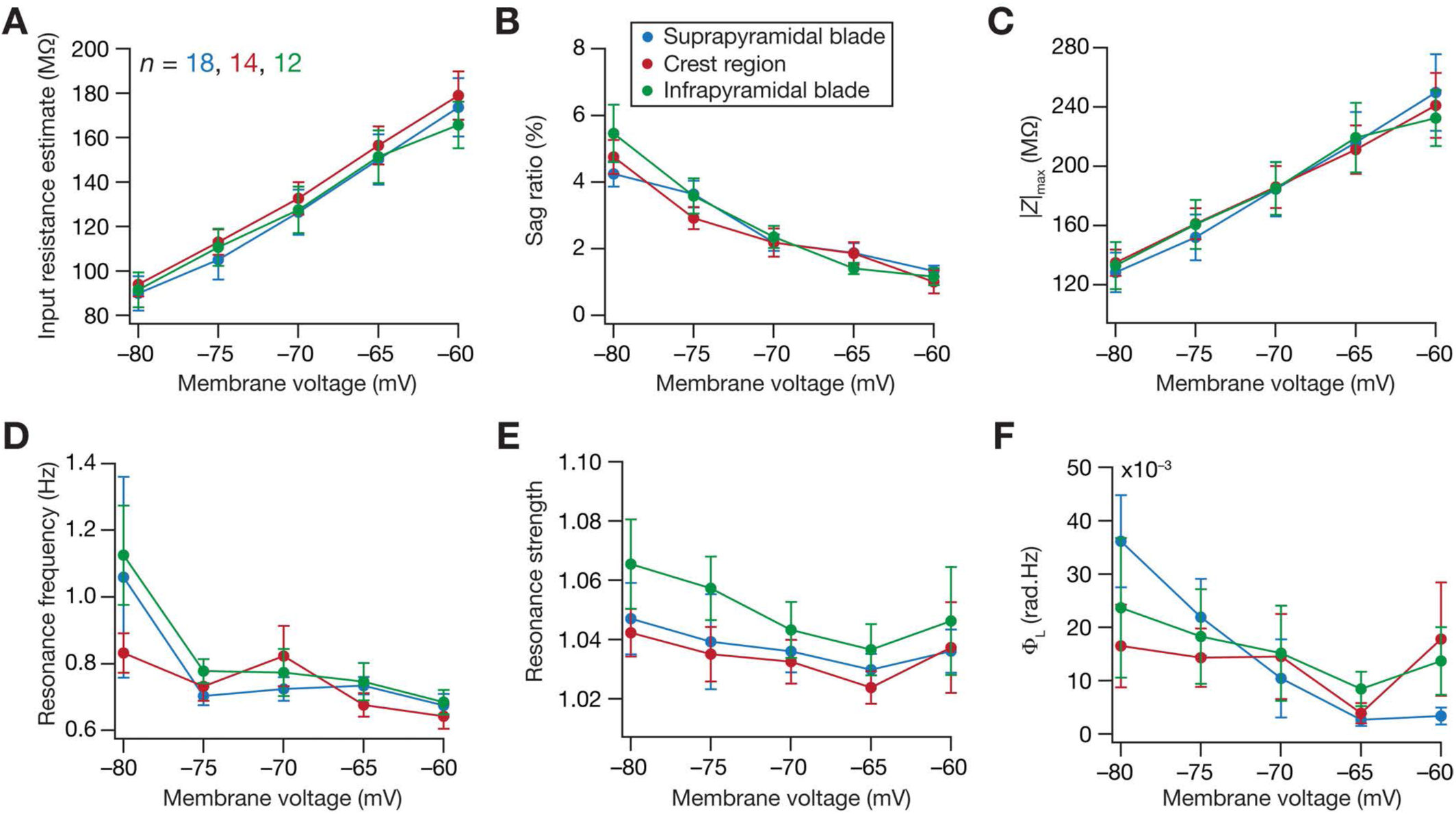
Voltage-dependence of subthreshold response properties of granule cells across the blades of the dentate gyrus. *A–F*: Voltage dependence of steady-state (*A–B*) and frequency-dependent (*C–F*) subthreshold measurements from the three DG sectors. The color-code for the three sectors are the same as those in Figure 2.

**Table 1.**
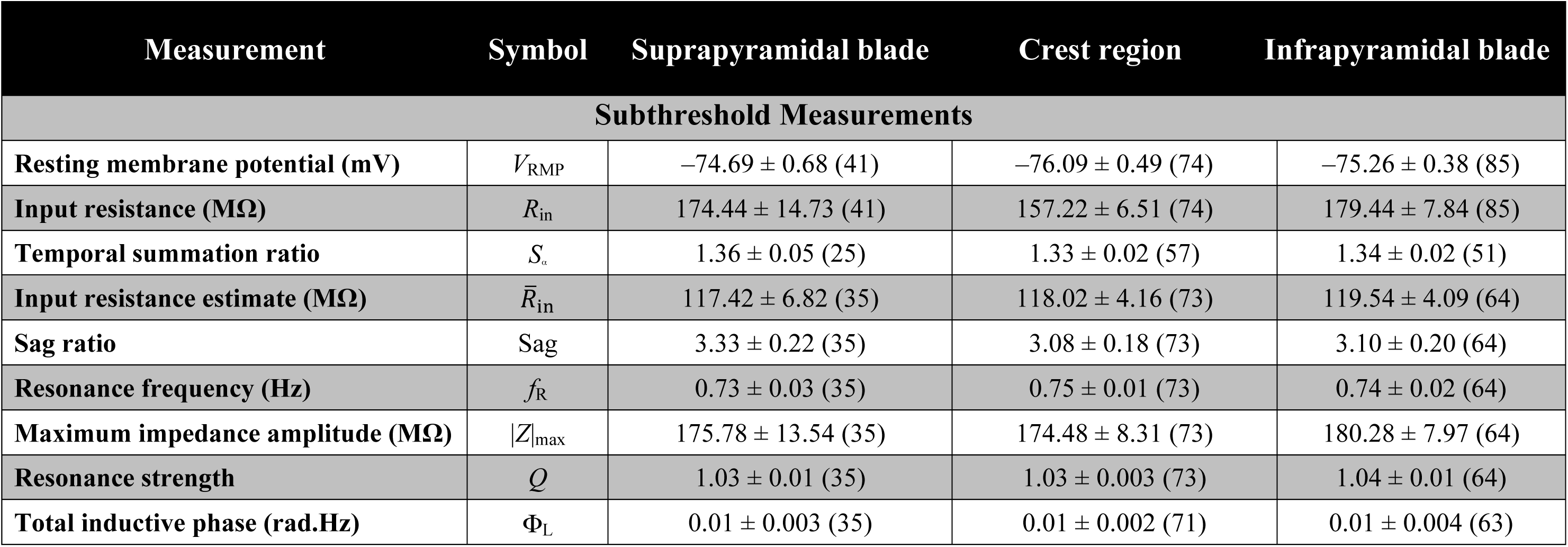
Subthreshold measurements when the respective current stimuli were injected with the cell resting at *V*_RMP_. Measurements are reported as mean ± SEM (*n*).

### Supra-threshold measurements

Action potential (AP) firing frequency was computed by extrapolating the number of spikes obtained during a 700 ms current injection to 1 s (Fig. 4*A*). Current amplitude of these pulse-current injections was varied from 0 pA to 250 pA in steps of 50 pA, to construct the firing frequency *vs.* injected current (*f–I*) plot (Fig. 4*B*). Various action potential related measurements (Malik et al. 2016; Mishra and Narayanan 2019) were derived from the voltage response of the cell to a 250 pA pulse-current injection (Fig. 4*C–D*). AP amplitude (*V*_AP_) was computed as the difference between the peak voltage of the spike (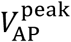) and *V*_RMP_ (Fig. 4*C*). The temporal distance between the timing of the first spike and the time of current injection was defined as latency to first spike (*T*_1AP_; Fig. 4*C*). The duration between the first and the second spikes was defined as the first inter-spike interval (*T*_1ISI_; Fig. 4*C*). AP half-width (*T*_APHW_; Fig. 4*D*) was the temporal width measured at the half-maximal points of the AP peak with reference to *V*_RMP_. The maximum 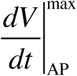 and minimum 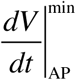 values of the AP temporal derivative were calculated from the temporal derivative of the action potential trace (Fig. 4*D*). The voltage in the AP trace corresponding to the time point at which the d*V*/d*t* crossed 20 V/s defined AP threshold (Fig. 4*D*). All supra-threshold measurements were obtained through current injections into the cell resting at *V*_RMP_, and were measured across each of the three sectors of the DG (Figs. 5–6).

**Figure 4.**
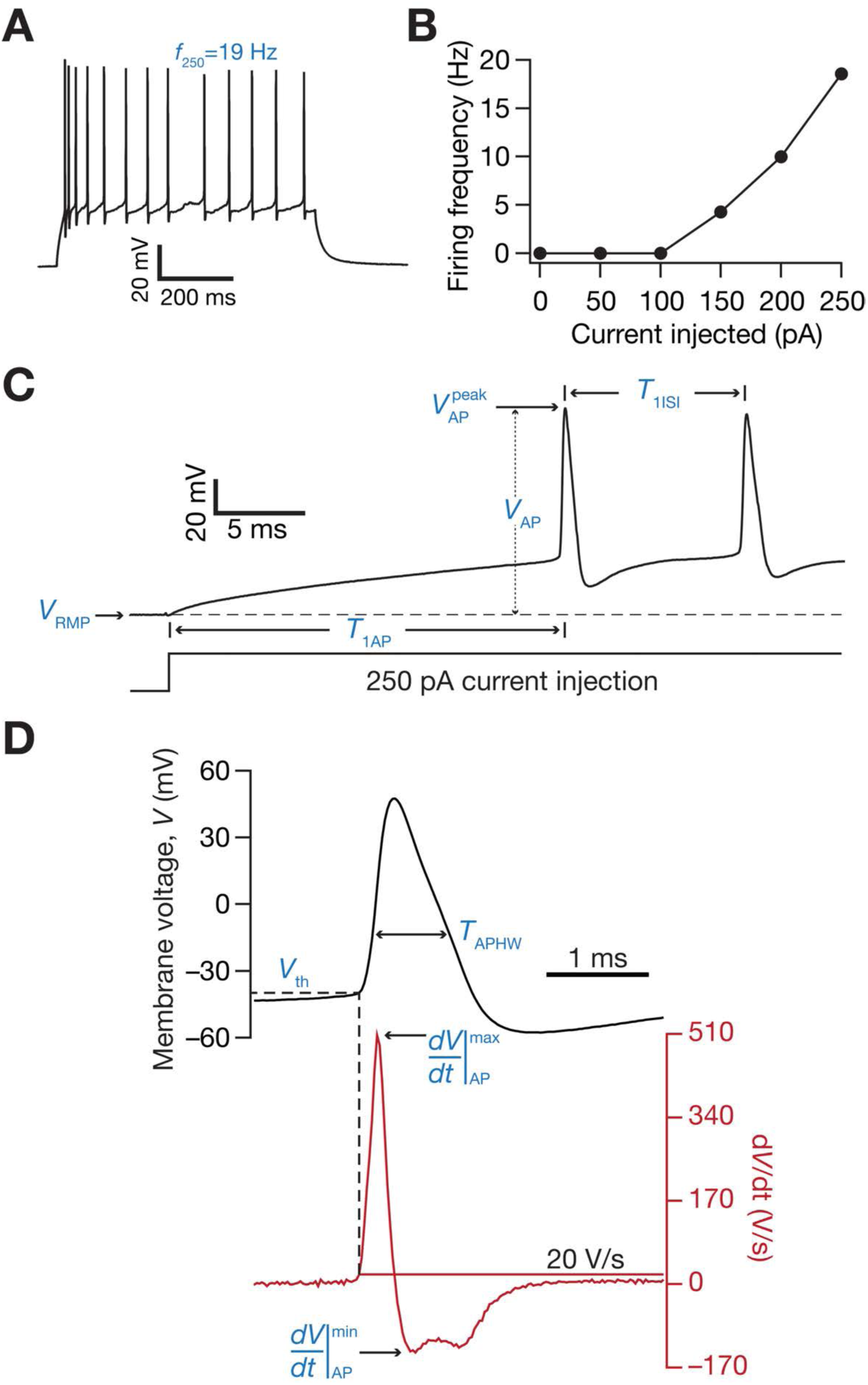
Electrophysiological protocols and measurements employed in the characterization of suprathreshold excitability of dentate gyrus granule cells. *A*: Voltage response (*left*) of the example neuron (same neuron as Figure 1) to a 700-ms current pulse of 250 pA. *B*: Frequency of firing plotted as a function of injected current amplitude for the example cell shown; note that these are firing frequencies converted from the number of spikes for a 700-ms duration. *C*: Zoomed version of the trace shown in panel *A*, illustrating electrophysiological measurements: *V*_RMP_ depicts resting membrane potential; *T*_1AP_ is the latency to the first action potential, measured from the time where the current injection was initiated; 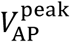 is the maximum voltage value measured on the first action potential; *V*_AP_ is the action potential amplitude, measured as the difference between 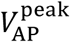 and *V*_RMP_; and *T*_1ISI_ is the first interspike interval measured as the temporal distance between the first and the second action potentials. *D*: Further zoomed version of the trace in panel *A* (black), along with its temporal derivative (d*V*/d*t*; red) illustrating electrophysiological measurements: the maximum 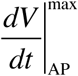 and minimum 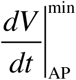 values of the action potential temporal derivative are depicted; the value on the voltage trace at the time when the value of the action potential temporal derivative crosses 20 V/s was assigned as the action potential threshold voltage (*V*_th_); the full width at half maximum of the action potential (with the maximum given by *V*_AP_) was assigned *T*_APHW_.

**Figure 5.**
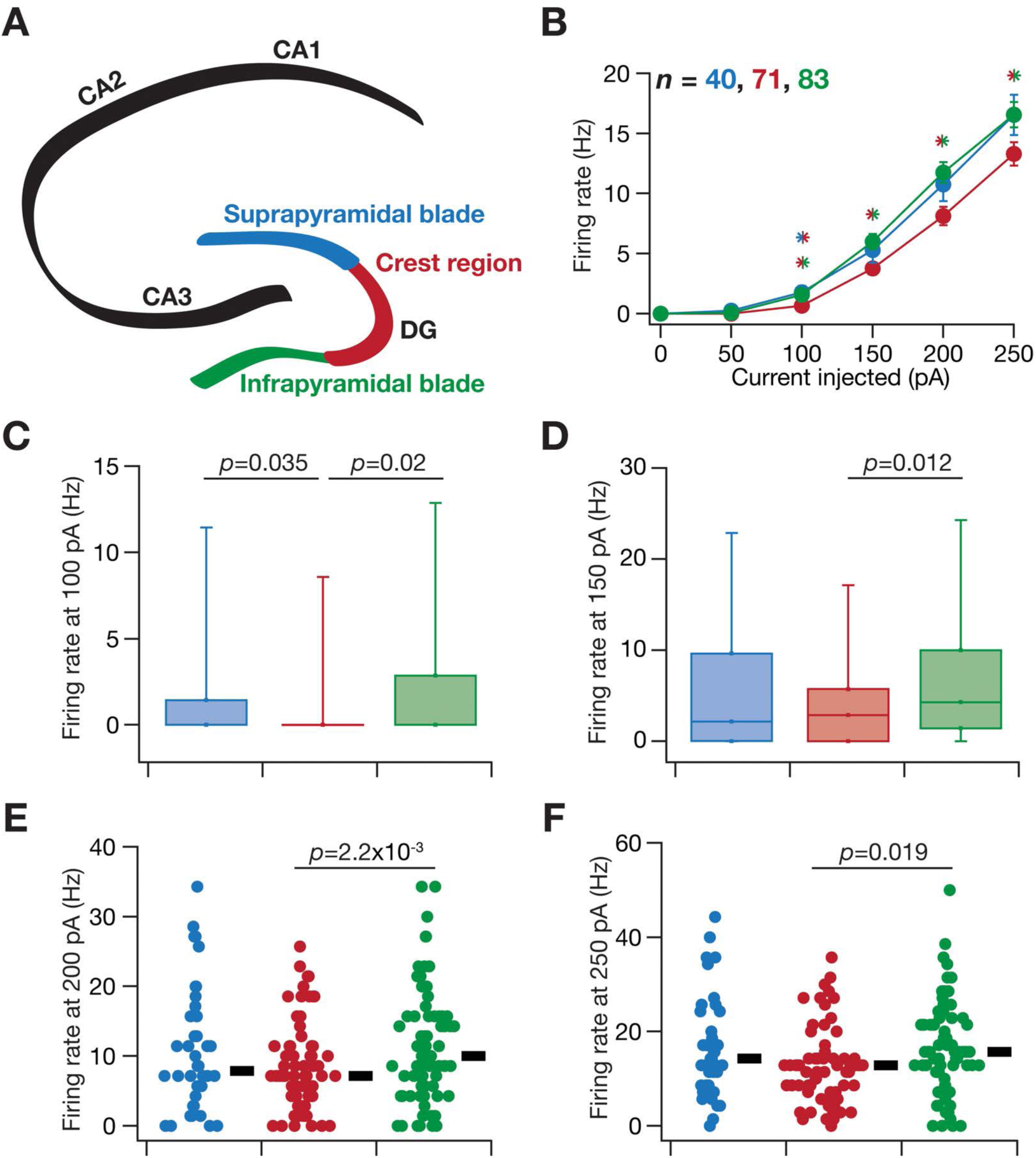
Heterogeneity in action potential firing frequency of granule cells across the blades of the dentate gyrus. *A*: Schematic of the hippocampus proper, showing the different CA (cornu ammonis) subregions (CA1, CA2 and CA3) and the three sectors of the dentate gyrus (infrapyramidal blade, crest region and suprapyramidal blade). The color codes associated with the three DG sectors apply for panels *B*–*F*. *B*: Frequency of firing plotted as functions of injected current amplitude for the populations of granule cells belonging to the three sectors. *: *p*<0.05, Student’s *t* test. The two colors in the asterisk represent the two populations across where significant differences were observed. *C*–*F*: Box plots depicting the heterogeneous action potential firing frequency of granule cells from the three DG sectors, for current injections of amplitude 100 pA (*C*) and 150 pA (*D*). Box plots are employed here because a significant proportion of cells did not fire action potentials, and representation with beeswarm plots exhibited clutters. *E*–*F*: Beeswarm plots depicting the heterogeneous action potential firing frequency of granule cells from the three DG sectors, for current injections of amplitude 200 pA (*E*) and 250 pA (*F*). None of the cells fired spontaneously and very few cells fired with 50 pA current injection. The black rectangles to the right of each beeswarm plot represent the median for the specified population. All measurements depicted in this figure were obtained through current injections into a cell resting at *V*_RMP_. The *p* values correspond to Wilcoxon rank sum test; *p* values less than 0.05 are shown.

**Figure 6.**
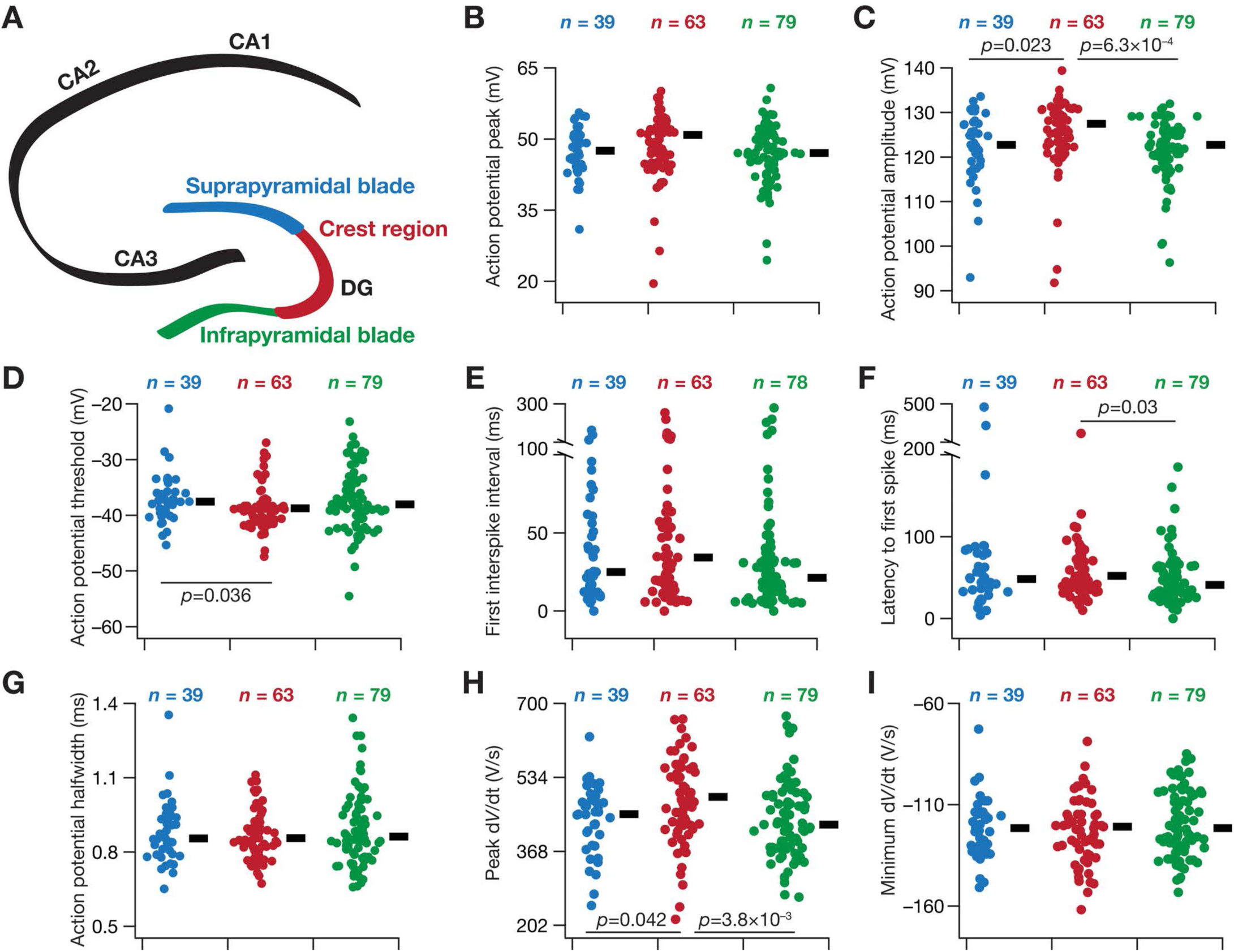
Heterogeneity in action potential properties of granule cells across the blades of the dentate gyrus. *A*: Schematic of the hippocampus proper, showing the different CA (cornu ammonis) subregions (CA1, CA2 and CA3) and the three sectors of the dentate gyrus (infrapyramidal blade, crest region and suprapyramidal blade). The color codes associated with the three DG sectors apply for panels *B*–*I*. *B*–*I*: Beeswarm plots depicting the heterogeneous action potential measurements from the three DG sectors. The black rectangles to the right of each beeswarm plot represent the median for the specified population. All measurements depicted in this figure were obtained through current injections into a cell resting at *V*_RMP_. The *p* values correspond to Wilcoxon rank sum test; *p* values less than 0.05 are shown.

### Pair-wise correlations between measurements

To assess pairwise relationship in 18 different sub- and supra-threshold measurements, we analyzed the scatter plot matrices of these measurements individually from each of the three sectors (Fig. 7*A–C*), as well of pooled measurements for all three sectors (Fig. 7*D*). We computed Pearson’s correlation coefficients for each of these pair-wise scatter plots and analyzed the distribution of correlation coefficients for each population (Fig. 7).

**Figure 7.**
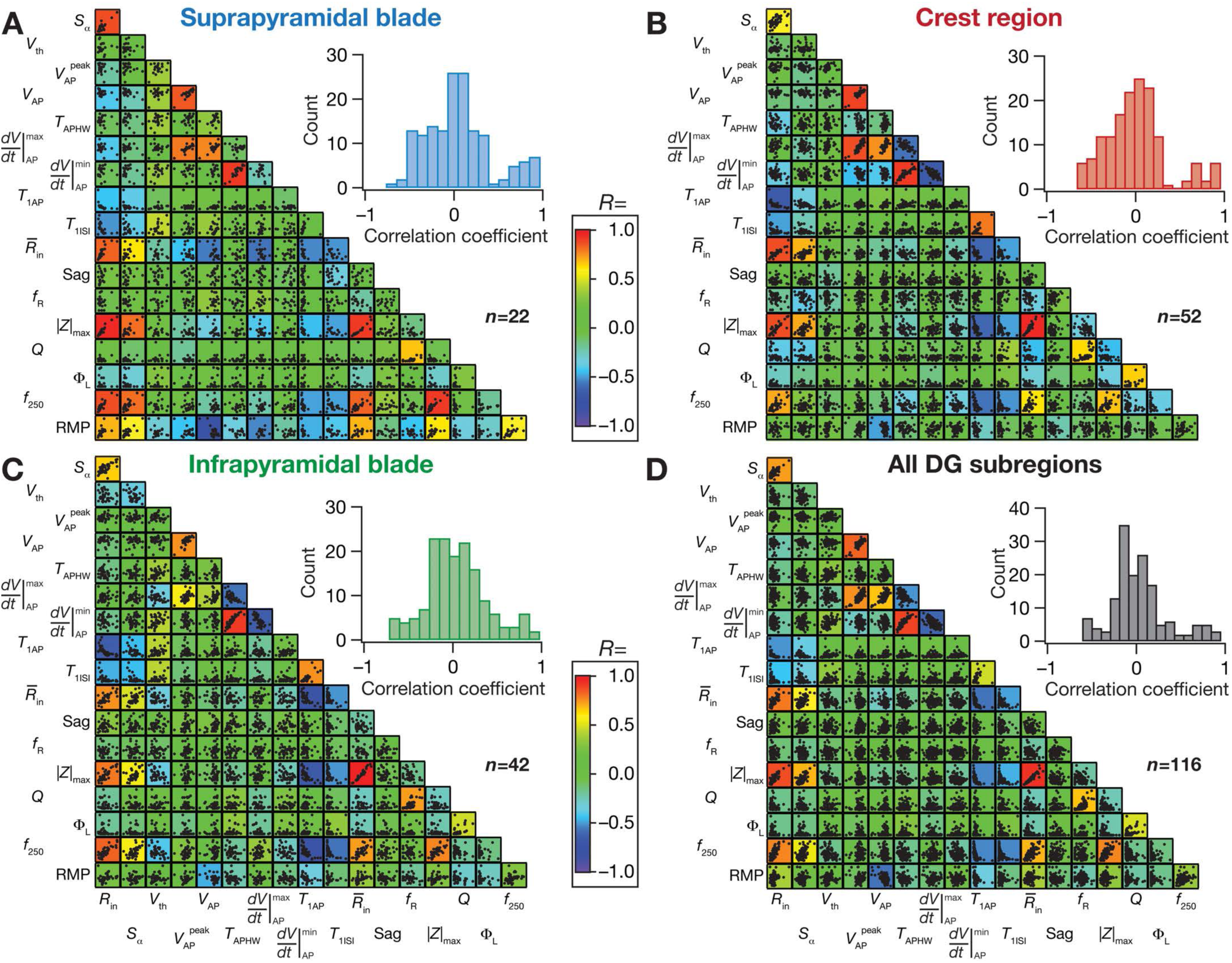
Differential correlations between sub- and supra-threshold measurements of granule cells across the blades of the dentate gyrus. *A–D*: Pair-wise scatter plot matrices of 18 sub- and supra-threshold measurements of granule cells recorded from the suprapyramidal blade (*A*), crest region (*B*), the infrapyramidal blade (*C*) and from all these sectors pooled together (*D*). These scatter plot matrices are overlaid on the corresponding color-coded correlation matrices. The inset in each panel represents the histograms of the correlation coefficients that are depicted by the correlation matrix. All measurements depicted in this figure were obtained through current injections into a cell resting at *V*_RMP_.

All data acquisition and analyses were performed using custom-written software in Igor Pro (Wavemetrics), and statistical analyses were performed using the R computing package (http://www.r-project.org/).

## RESULTS

In assessing the intrinsic response properties of DG granule cells across its different blades, we performed patch-clamp electrophysiological recordings under current clamp mode at the cell body of visually identified granule cells. We first characterized the response properties of these neurons using a range of subthreshold electrophysiological measurements (Fig. 1). We measured input resistance from the steady-state voltage response to pulse current injections of different depolarizing and hyperpolarizing amplitude (Fig. 1*A*). Input resistance, a steady-state measure of neuronal gain and excitability, falls inadequate in characterizing neuronal response properties to ethologically relevant time-varying signals. Therefore, to assess neuronal response properties to time-varying signals, we injected multiple current stimuli that mimicked excitatory postsynaptic currents (EPSC) into neurons to understand temporal summation properties (Magee 1998; 1999). We found considerable temporal summation of EPSPs, with the value of temporal summation ratio above unity (Fig. 1*B*).

As the dentate gyrus resides in an oscillatory neural network (Bland 1986; Buzsaki 2002; Colgin 2013; 2016; Sainsbury and Bland 1981; Winson 1978; 1974), we assessed neuronal response properties to sinusoidal stimulus of different frequencies. A standard stimulus that is employed in assessing frequency-dependent response properties is the chirp current stimulus (Fig. 1*C*), which is a sinusoidal current of constant amplitude but with monotonically increasing frequency (Gimbarzevsky et al. 1984; Hutcheon and Yarom 2000; Krueppel et al. 2011; Stegen et al. 2012). We found that the frequency-dependent response profile was low-pass in nature for all measured subthreshold voltages (Figs. 1*D–E*), in stark contrast to the band-pass profiles of other cell types within the hippocampal formation, such as the medial entrorhinal stellate neurons (Erchova et al. 2004; Giocomo et al. 2007) and CA1 pyramidal neurons (Hu et al. 2009; Hu et al. 2002; Narayanan and Johnston 2008; 2007; Pike et al. 2000). The gain of the system, measured as impedance amplitude, reduced especially across lower frequencies with hyperpolarization of membrane potential (Fig. 1*D–E*), and reflected in the maximal impedance amplitude |*Z*|_max_ reducing with hyperpolarization (Fig. 1*E*).

The advantage of employing impedance as an excitability measure is two-fold. First, impedance amplitude measures neuronal excitability as a function of input frequency providing a frequency-dependent excitability metric, and second, impedance phase provides the temporal relationship between the voltage response and current input at various input frequencies (Mauro 1961; Mauro et al. 1970; Narayanan and Johnston 2008; Sabah and Leibovic 1969). We computed the impedance phase at all measured frequencies, and found the voltage response to lag the injected current at all membrane voltages (Fig. 1*F*). This is in striking contrast with CA1 pyramidal and entorhinal stellate cells, where the impedance phase profile is biphasic, with the voltage-response leading the current injection at theta-frequency range, and lagging beyond theta-frequency range (Erchova et al. 2004; Mittal and Narayanan 2018; Narayanan and Johnston 2008). We also noticed that the maximal lag observed, at the highest measured frequency (15 Hz) reduced with increasing hyperpolarization, pointing to a systematic increase in inductive component. This increased inductive component, however, was unable to drive impedance phase values to positive ranges even in the most hyperpolarized recordings. We quantified this lack of positive impedance phase values as Φ_L_, the area under the inductive (positive) part of the ZPP, which was zero at all voltages where chirp responses were recorded (Fig. 1*F*).

As functions of membrane voltage, both input resistance as well as maximal impedance amplitude reduced with hyperpolarization, indicating a reduction in overall neuronal excitability at hyperpolarized voltages (Fig. 1*G*). The resonance frequency (*f*_R_) was less than 1 Hz and the resonance strength was close to unity at all measured voltages (Fig. 1*H*), indicating low-pass response characteristics of this dentate granule cell.

### Heterogeneities in subthreshold measurements across different blades of the dentate gyrus

How do these different steady-state and frequency-dependent subthreshold measures of granule cell response vary across the different blades of the dentate gyrus? To address this question, we measured the different electrophysiological properties (Fig. 1) at resting membrane potential from granule cells located within the three prominent sectors within the DG: the suprapyramidal blade, the crest region and the infrapyramidal blade (Fig. 2*A*). First, we found considerable cell-to-cell variability in each of these response properties, spanning all the three sectors (Fig. 2*B–I*; Table 1). For instance, whereas the median value of input resistance was around 150 MΩ across all three sectors, the input resistance spanned a large range from tens to hundreds of MΩ (Fig. 2*C*). Second, we found that none of these subthreshold measurements were significantly different across the three sectors (Fig. 2*B–I*) implying the similarity in the degree of heterogeneity across all sectors of DG.

These measurements confirmed that DG granule cells across all three sub-structures lack prominent sag (Fig. 2*E*) that is characteristic of the expression of resonance. This was consistent with the resonance frequency of these neurons falling around 1 Hz (Fig. 2*H*), with the resonance strength centered on unity (Fig. 2*I*). Together, these measurements indicate that DG granule cells across all three sectors exhibited low-pass response characteristics. The temporal summation of alpha current inputs showed the fifth EPSP to have higher amplitude than the first (temporal summation ratio > 1) for most recorded neurons across all three sectors, indicating an enhanced temporal summation in these neurons (Fig. 2*F*). However, cells within each sector exhibited significant heterogeneity in terms of how they responded to the train of alpha current inputs, with temporal summation ranging from a value just lower than unity to values greater than 1.5 in certain cells (Fig. 2*F*).

### Dentate gyrus granule cells exhibited low-pass response characteristics and lacked inductive lead in their impedance phase profiles at all subthreshold voltages and across different blades

How does neuronal excitability change as a function of membrane voltage? Do these cells exhibit band-pass characteristics at more depolarized or hyperpolarized voltages, similar to the voltage-dependent resonance properties observed in CA1 pyramidal and entorhinal stellate neurons? Are there sector-specific differences in terms of how neurons respond to different voltages? To address these questions, we altered the neuronal membrane potential employing DC current injection, and recorded subthreshold measurements at five different voltage values (Fig. 3). We found neuronal excitability to reduce with increased hyperpolarization, inferred from hyperpolarization-induced reductions in input resistance (Fig. 3*A*) and maximum impedance amplitude (Fig. 3*C*) across all three sectors. Although there was a small increase in sag (Fig. 3*B*) and resonance frequency (Fig. 3*D*) at hyperpolarized voltages, these increases were not as prominent as CA1 pyramidal neurons or entorhinal stellate neurons (Erchova et al. 2004; Hu et al. 2009; Hu et al. 2002; Narayanan and Johnston 2007), and the resonance strength continued to center at unity (Fig. 3*E*). Furthermore, the total inductive phase was negligibly small across all voltages, confirming the absence of an inductive phase lead in the impedance profile of granule cells across all three sectors (Fig. 3*F*). Together these results demonstrated that DG granule cells exhibit low-pass response properties with a distinct absence of inductive lead in the impedance phase, at all subthreshold voltages and across the three sectors of the dentate gyrus.

### Heterogeneities in firing properties and action potential measurements across different blades of the dentate gyrus

How do neuronal firing profiles and action potential properties vary across the different sectors of the dentate gyrus? Are these suprathreshold measurements heterogeneous within each sector? We injected depolarizing current pulses of different amplitudes to assess the firing profile (Fig. 4*A–B*) and several metrics associated with action potentials (Fig. 4*C–D*; Table 2) of DG granule cells. Across the three DG sectors, the firing profile of dentate granule cells (Fig. 4*B*, Fig. 5*B–F*) reflected class I excitability, where the neuron was capable of eliciting firing at arbitrarily low frequencies (Hodgkin 1948; Ratte et al. 2013). In addition, beyond rheobase current (which was between 50–150 pA in most recorded neurons; Fig. 5*B–D*), the firing rate increase as a function of injected current was fairly linear (Fig. 5*B*). These observations, along with the low-pass response characteristics and positive temporal summation observed earlier (Figs. 1–3), pointed to the DG granule neurons acting as integrators of incoming information (Ratte et al. 2013).

**Table 2.**
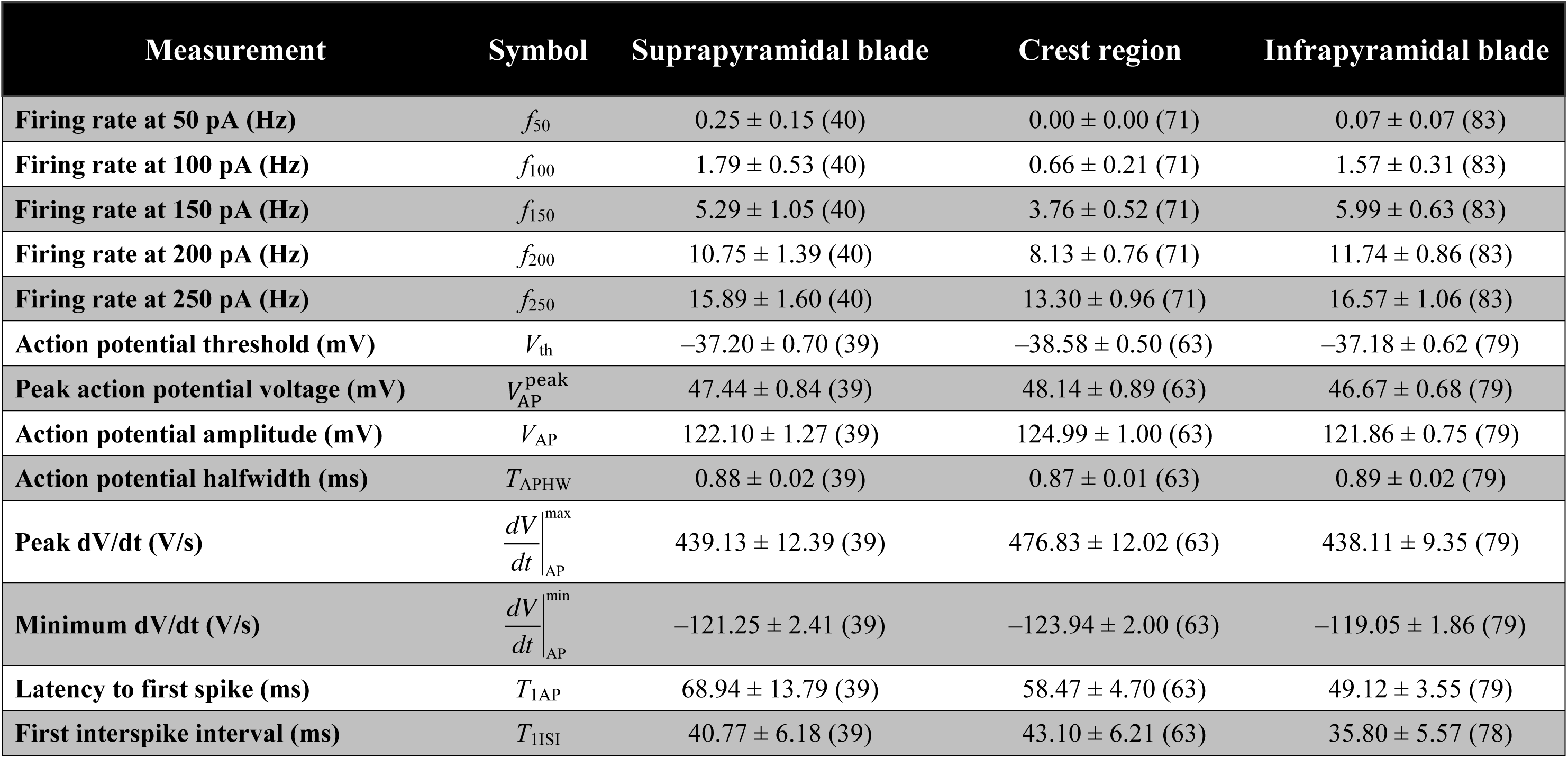
Suprathreshold measurements when the respective current stimuli were injected with the cell resting at *V*_RMP_. Measurements are reported as mean ± SEM (*n*).

We also observed considerable cell-to-cell variability in firing frequencies within each of the three sectors. For instance, for a pulse current injection of 250 pA, the median firing rate of these neurons was around 15 Hz across all three sectors (Fig. 5*F*). However, this rate varied over a large range spanning 0 (no spikes) for certain cells to ∼40 Hz in certain others. Thus, the mean (Fig. 5*B*) or median (Fig. 5*F*) firing rate values should be treated with caution, because there are cells that do not spike even for 250 pA current injection, whereas certain others spike with rates more than twice the mean/median value. This heterogeneity was observed across all current injections that were assessed (Fig. 5*C–F*). Importantly, although there was not a large difference in overall range of firing rates observed across the three sectors (Fig. 5*C–F*), granule cells from the crest region showed significantly (*p*<0.05, Student’s *t* test in Fig. 5C and Wilcoxon rank sum test in Figs. 5*D–F*) reduced excitability compared to cells in the infrapyramidal blade across all measured current injections (Fig. 5*B–F*).

We next analyzed individual spikes from granule cells recorded from each of the three sectors and derived metrics that quantified spike threshold, width, amplitude, depolarizing and repolarizing kinetics. Similar to our observations with subthreshold properties (Fig. 2) and action potential firing properties (Fig. 5), we found significant cell-to-cell variability in these measurements even within a given sector (Fig. 6*B–I*). Across populations, we found action potential amplitude to be significantly lower in granule cells from the crest region compared to those from the two blades (Fig. 6*C*). Action potential threshold was significantly hyperpolarized in crest region cells compared to those in the suprapyramidal blade (Fig. 6*D*). The latency to first spike was significantly lower in the infrapyramidal population of granule cells compared to cells in the crest region (Fig. 6*F*). Finally the peak derivative of the action potential trace was significantly higher for cells in the crest region, compared to cells in the two blades (Fig. 6*H*). We noted that although these differences were statistically significant, the range of values of these measurements from neurons of all three sectors was not very different (Fig. 6*B–I*). In addition, as the differences in median (Fig. 6) or mean values (Table 2) of these measurements across the sectors were not large (as a fraction of the respective range of each measurement), it might be infeasible to infer large differences in action potential properties across DG sectors based on these measurements (especially in light of the enormous heterogeneity in each measurement, represented in Fig. 6 and quantified as SEM in Table 2).

### A large proportion of sub- and supra-threshold measurements from all blades of the dentate gyrus exhibited weak pairwise correlations

Are the different sub- and supra-threshold measurements correlated? Are these correlations distinct across the different blades of the dentate gyrus? Correlations in measurements provide clues about relationships between sub- and supra-threshold measurements, apart from pointing to the possibility of similar ion channels mechanisms underlying these distinct measurements. We plotted pairwise scatter plots of these intrinsic measurements from granule cells recorded from the suprapyramidal blade (Fig. 7*A*), the crest region (Fig. 7*B*), the infrapyramidal blade (Fig. 7*C*) and the pooled population containing cells from all sectors (Fig. 7*D*). We computed Pearson’s correlation coefficient for each of these pairwise scatter plots (Fig. 7*A–D*). Broadly, we found the correlation matrices across the four populations to be fairly similar, with a large number of measurement pairs showing weak pairwise correlations (between –0.5 to +0.5; see inset histograms in Fig. 7*A–D*). A small set of pairs showed strong positive or negative correlations, and this set was also broadly common across the 4 matrices (Fig. 7*A–D*). Among measurements that exhibited strong positive correlations were the different measures of excitability (*R*_in_, 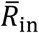, |*Z*|_max_, *f*_250_), the measures that were reflective of the depolarizing (*V*_AP_, 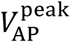 and 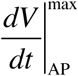) or repolarizing (*T*_APHW_ and 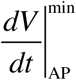) phase of action potential. Interestingly, temporal summation ratio also showed strong positive correlations with the different measures of excitability. Among measurements that exhibited strong negative correlations were the latency to first spike *vs.* each measure of excitability and the first ISI *vs.* each measure of excitability. Thus, except for a small subset of highly correlated measurements that are derived from common physiological substrates, the large subset of uncorrelated measurements suggest that the set of measurements employed here in characterizing DG granule cells are assessing distinct aspects of their physiology.

## DISCUSSION

We performed whole-cell patch-clamp electrophysiological recordings of granule cells from the three sectors of the rat DG, and systematically measured their sub- and supra-threshold electrophysiological characteristics. Our recordings demonstrate that the granule cells within the three different DG sectors manifest considerable heterogeneities in their intrinsic excitability, temporal summation, action potential characteristics and frequency-dependent response properties. Assessing neuronal responses to time-varying inputs, we find that DG neurons, across all three sectors, showed positive temporal summation of their responses to current injections that mimicked excitatory postsynaptic currents. Next, we demonstrated that the impedance amplitude profile manifested low-pass characteristics and the impedance phase profile distinctly lacked positive phase values at all measured frequencies, voltages and for all DG sectors. Consistent with the lack of an inductive component in either the impedance amplitude or phase profiles, neurons across all sectors showed little to no sag in their voltage responses to hyperpolarizing or depolarizing pulse current injections. In what follows, we explore some implications for the expression of heterogeneities inherent to these individual sectors, and for the absence of inductive components in the impedance profiles.

### Impact of heterogeneities on information processing in the dentate gyrus

The impact of heterogeneities on neural physiology and information processing is well established (Angelo et al. 2012; Anirudhan and Narayanan 2015; Cadwell et al. 2016; Chelaru and Dragoi 2008; Das et al. 2017; Ecker et al. 2011; Fuzik et al. 2016; Gjorgjieva et al. 2016; Goaillard et al. 2009; Grashow et al. 2010; Kohn et al. 2016; Marder 2011; Marder and Goaillard 2006; Marder et al. 2014; Marder and Taylor 2011; Mishra and Narayanan 2019; Mittal and Narayanan 2018; Mukunda and Narayanan 2017; Nusser 2009; Padmanabhan and Urban 2010; Prinz et al. 2004; Rathour and Narayanan 2019; 2014; 2012a; Renart et al. 2003; Shamir and Sompolinsky 2006; Srikanth and Narayanan 2015; Tikidji-Hamburyan et al. 2015; Tripathy et al. 2013; Voliotis et al. 2014; Wang and Buzsaki 1996; Zhou et al. 2013). Within the context of dentate gyrus physiology, what could be the impact of the heterogeneities observed in their sub-(Fig. 2) and supra-threshold (Fig. 5) excitability properties? The dentate gyrus has been implicated in pattern separation and response decorrelation. There are lines of evidence from different brain regions that heterogeneities in intrinsic neuronal properties could play a critical role in effectuating such response decorrelation (Mishra and Narayanan 2019; Padmanabhan and Urban 2010; Tripathy et al. 2013). The heterogeneities characterized here could contribute to channel and pattern decorrelation in the DG network.

The precise shape of synaptic plasticity profiles in neurons is critically reliant on the local and global excitability properties of the postsynaptic neurons (Anirudhan and Narayanan 2015; Golding et al. 2002; Johnston et al. 2003; Johnston and Narayanan 2008; Narayanan and Johnston 2010; Sehgal et al. 2013; Sjostrom and Hausser 2006; Watanabe et al. 2002). This reliance is not dependent only on suprathreshold conductances that enable spike generation and sustain backpropagation of action potentials and dendritic spikes, but also on subthreshold conductances that regulate synaptic potentials and their temporal summation (Anirudhan and Narayanan 2015; Chen et al. 2006; Malik and Johnston 2017; Narayanan and Johnston 2010; Nolan et al. 2004). Therefore, the heterogeneities reported here in temporal summation, apart from sub- and supra-threshold excitability properties could translate to differences in synaptic plasticity profiles of these neurons. Such differential impact of synaptic plasticity protocols, which could concomitantly and differentially alter neuronal intrinsic properties as well (Lopez-Rojas et al. 2016; Stegen et al. 2012; Titley et al. 2017), could also mediate the recruitment of specific cells in memory formation and in the efficacy of memory retrieval (Josselyn and Frankland 2018; Pignatelli et al. 2019; Silva et al. 2009; Yiu et al. 2014; Zhou et al. 2009). Specifically, the baseline differences in excitability reported here could, in conjunction with other mechanisms such as neuromodulation, enable the recruitment of specific cells during memory formation. Together, it is important that experimental interpretations and computational models account for these prominent heterogeneities in the physiological response properties of granule cells.

### Granule cells are integrators of afferent information: Class I excitability coupled with lack of sag, resonance and positive impedance phase

Hodgkin (Hodgkin 1948) had classified excitability into three distinct classes based on initiation of repetitive action potential firing through constant current injection. Axons that were capable of responding over a wide range of frequencies, especially at arbitrarily low frequencies were designated as class I. Axons that were classified as class II exhibited a pronounced supernormal phase, whereby the frequency of action potential firing was largely invariant to the injected current amplitude after the first spike was elicited (for currents beyond the rheobase current). Class III axons were those that elicited a second spike (beyond the first spike) only with difficulty or not at all (Hodgkin 1948). This classification has provided an invaluable tool to understand neuronal excitability, neural coding spike initiation dynamics, neuronal operating characteristics and phase resetting curves in a broadly unified, with the ionic mechanisms underlying these classes of excitability well understood (Das and Narayanan 2015; 2014; 2017; Das et al. 2017; Ermentrout 1996; Hodgkin 1948; Prescott et al. 2008a; Prescott et al. 2006; 2008b; Ratte et al. 2013). Specifically, it is now recognized that the different classes of excitability are consequent to cooperation or competition between fast inward currents and slow outward currents. Cooperation between these two classes of currents yield class I excitability, whereas competition yields class II/III excitability. Importantly, these classes of excitability have been linked to the ability of neurons to act as integrators (class I) or as coincidence detectors (class II/III), with the synergistic interactions among channels capable of sliding the operating mode of a neuron along the integrator–coincidence detector continuum (Das and Narayanan 2015; 2014; 2017; Das et al. 2017; Ratte et al. 2013).

Our results demonstrate that the *f–I* curve of DG granule cells are capable of firing at arbitrarily low frequencies, with firing frequency clearly dependent on input current injection (Fig. 5*B*), pointing to class I excitability characteristics. In addition, the absence of sag, resonance and positive impedance phase also point to absence of a dominant slow outward current that can contribute to class II/III excitability or coincidence detection capabilities (Das and Narayanan 2015; 2014; 2017; Das et al. 2017; Krueppel et al. 2011; Ratte et al. 2013). Together these results clearly point to dentate granule cell somata acting as integrators of afferent information (also see (Aimone et al. 2011)). However, it should be noted that operating modes of neurons could change in response to several factors, including activity-dependent plasticity of channels and receptors, neuromodulation and changes in afferent activity patterns (Das and Narayanan 2015; 2014; 2017; Das et al. 2017; Prescott et al. 2008a; Prescott et al. 2006; 2008b; Ratte et al. 2013). Future studies should therefore assess the spike-triggered average of DG neurons to understand their operating modes, the roles of different ion channels in regulating operating mode across their somato-dendritic axis and the information processing strategies employed by the DG neurons (Das and Narayanan 2015; 2014; 2017; Das et al. 2017; Krueppel et al. 2011; Schmidt-Hieber et al. 2007).

Despite the well-established expression of the hyperpolarization-activated cyclic nucleotide gated (HCN) channels in granule cells (Bender et al. 2003; Krueppel et al. 2011; Stegen et al. 2012; Surges et al. 2012), they don’t express impedance resonance or positive impedance phase (Figs. 1–3) unlike CA1 pyramidal neurons or entorhinal stellate neurons (Das et al. 2017; Erchova et al. 2004; Hu et al. 2009; Hu et al. 2002; Mittal and Narayanan 2018; Narayanan and Johnston 2008; 2007). However, it is established that the expression of impedance resonance is not just dependent on the expression of a resonating conductance, but on the density of the resonating conductance and the leak conductance in the neuron (Hutcheon et al. 1996a; b; Hutcheon and Yarom 2000; Narayanan and Johnston 2008; 2007; Rathour et al. 2016; Rathour and Narayanan 2014; 2012a; b; Zemankovics et al. 2010), on the time constants of the resonating conductance (Hutcheon et al. 1996a; Hutcheon and Yarom 2000; Rathour and Narayanan 2012a), on morphological properties (Dhupia et al. 2014), on the relative expression profiles of other subthreshold channels and interactions of these channels with the resonating conductances (Das et al. 2017; Rathour et al. 2016; Rathour and Narayanan 2019; 2014; 2012a; Stegen et al. 2012). The frequency-dependent response analyses presented in our study also provide indirect evidence for the expression of HCN channels, through the reduction of excitability at hyperpolarized voltages (Fig. 3), through a hyperpolarization-induced suppression of gain that is dominant at low frequencies (Fig. 1*E*) and through the reduction in the capacitive lag in the impedance profiles (Fig. 1*F*) with hyperpolarization.

### Future directions

Future studies should explore all of somato-dendritic, infrapyramidal-suprapyramidal, dorso-ventral, and superficial-deep axes of the dentate gyrus, and characterize intrinsic heterogeneities expressed not just in granule cells, but also in other cell types including the basket cells and the mossy cells. In addition, these studies should explore if there are systematic gradients in voltage-gated and ligand-gated channels across these different axes, which would alter the processing and encoding strategies associated with these neurons. The assessment of these heterogeneities and gradients are especially essential, given the gradients that are observed in intrinsic and synaptic properties of other spatially proximal cell types, including the CA1 pyramidal neurons and entorhinal stellate cells (Cembrowski et al. 2016; Cembrowski and Spruston 2019; Danielson et al. 2016; Dougherty et al. 2012; Dougherty et al. 2013; Giocomo and Hasselmo 2009; Giocomo et al. 2007; Kjelstrup et al. 2008; Lee et al. 2014; Malik et al. 2016; Marcelin et al. 2012; Maroso et al. 2016; Mizuseki et al. 2011; Soltesz and Losonczy 2018; Strange et al. 2014; Valero et al. 2015). Our analyses also emphasize that experimental interpretations and computational models should not extrapolate DG cell excitability from summary statistics or treat DG neurons as a homogenous population, but should account for the extensive heterogeneities prevalent in the physiological response properties of granule cells.

## Acknowledgments

This work was supported by the Wellcome Trust-DBT India Alliance (Senior fellowship to RN; IA/S/16/2/502727), the Department of Biotechnology through the DBT-IISc partnership program (RN), the Revati & Satya Nadham Atluri Chair at IISc (RN) and the Ministry of Human Resource Development (RN & PM). The authors thank Divyansh Mittal for helpful discussions and for comments on a draft of this manuscript.

## Notes

**Competing financial interests** The authors declare no conflict of interest.

